# Fluid flow reconstruction around a free-swimming sperm in 3D

**DOI:** 10.1101/2024.05.29.596379

**Authors:** Xiaomeng Ren, Paul Hernández-Herrera, Fernando Montoya, Alberto Darszon, Gabriel Corkidi, Hermes Bloomfield-Gadêlha

**Author notes:** **Author for correspondence:** Hermes Bloomfield-Gadelha,.

## Abstract

We investigate the dynamics and hydrodynamics of a human spermatozoa swimming freely in 3D. We simultaneously track the sperm flagellum and the sperm head orientation in the laboratory frame of reference via high-speed high-resolution 4D (3D+t) microscopy, and extract the flagellar waveform relative to the body frame of reference, as seen from a frame of reference that translates and rotates with the sperm in 3D. Numerical fluid flow reconstructions of sperm motility are performed utilizing the experimental 3D waveforms, with excellent accordance between predicted and observed 3D sperm kinematics. The reconstruction accuracy is validated by directly comparing the three linear and three angular sperm velocities with experimental measurements. Our microhydrodynamic analysis reveals a novel fluid flow pattern, characterized by a pair of vortices that circulate in opposition to each other along the sperm cell. Finally, we show that the observed sperm counter-vortices are not unique to the experimental beat, and can be reproduced by idealised waveform models, thus suggesting a fundamental flow structure for free-swimming sperm propelled by a 3D beating flagellum.

## 1. Introduction

Fluids are ubiquitous in gamete and embryo transport, serving as the fundamental basis for microorganism locomotion [1–5]. Experimental research on the fluid flow around swimming microorganisms generally falls into two categories: direct measurements of the flow field using particle image velocimetry (PIV) [6–8] and theoretical reconstructions of the flow field based on experimentally measured beat patterns [9–11]. Experimental analysis of collective motion in fresh semen was performed by [6], revealing quasi-2D turbulence in this concentrated active suspension through PIV. Similarly, [8] visualized fluid flow around a tethered human sperm using PIV, uncovering an average spiral flow structure in the flagellum rolling plane. In contrast, [11] extracted the planar beating pattern of a human sperm via digital imaging microscopy, and predicted both the flow around the cell and its trajectory based on the experimental waveform. No study to date attempted to reconstruct the fluid flow pattern around a free-swimming sperm powered by a 3D beating flagellum.

In this work, we track the 3D free-swimming motion of human spermatozoa by imaging microscopy, unveiling the experimental waveform as seen from the body frame of reference, relative to its head. We reconstruct the observed sperm movements numerically using the experimental waveform, and confirm our reconstructions through qualitative and quantitative comparisons with experimental data. Remarkably, we report an excellent agreement between observed and predicted sperm kinematics (3D cell trajectories and speeds), validating both the empirical method and the micro-hydrodynamic theory commonly employed for microorganism propulsion. We proceed to investigate the hydrodynamics surrounding the human sperm, and report a novel fluid flow pattern currently unreported, characterized by a pair of vortices that circulates in opposition around the sperm body in 3D. This flow structure departs from previous reports, only known for non-swimming or 2D beating sperm [6, 11–13]. For instance, [14] elucidates 2D swirling flows within the flagellum beating plane, while [15] presents vortex coordination among self-organized cells as a result of fluid interaction. Finally, we show that this flow pattern can be replicated by idealised waveform models, suggesting the counter-vortices a fundamental flow structure for 3D free-swimming sperm.

## 2. Results

### 2.1 3D human sperm motility reconstruction

We track the 3D free swimming of a human sperm in the laboratory frame of reference (lab frame), as previously reported in [16], the motility of which is shown in Fig. 1(b) and Supplemental Material, Video 1. Viewed from head to tail, the sperm head spins counterclockwise (CCW) around its longitudinal axis, and the trajectory of the mid-flagellar point is characterized with local CCW loops and global clock-wise (CW) revolutions, as described by [17, 18]. By detecting the head position, ***X***_*l*_ = (*x*_*l*_, *y*_*l*_, *z*_*l*_), and 3D head orientation, ***E*** = [***e***_1_, ***e***_2_, ***e***_3_], in the lab frame, we obtain the 3D sperm waveform relative to the body frame of reference independent of translation and rotation, details see Methods. Fig. 1(a) and Supplemental Material, Video 2 display the 3D flagellar beating in the body frame, where the head appears is stationary relative to the flagellum.

**Figure 1.**
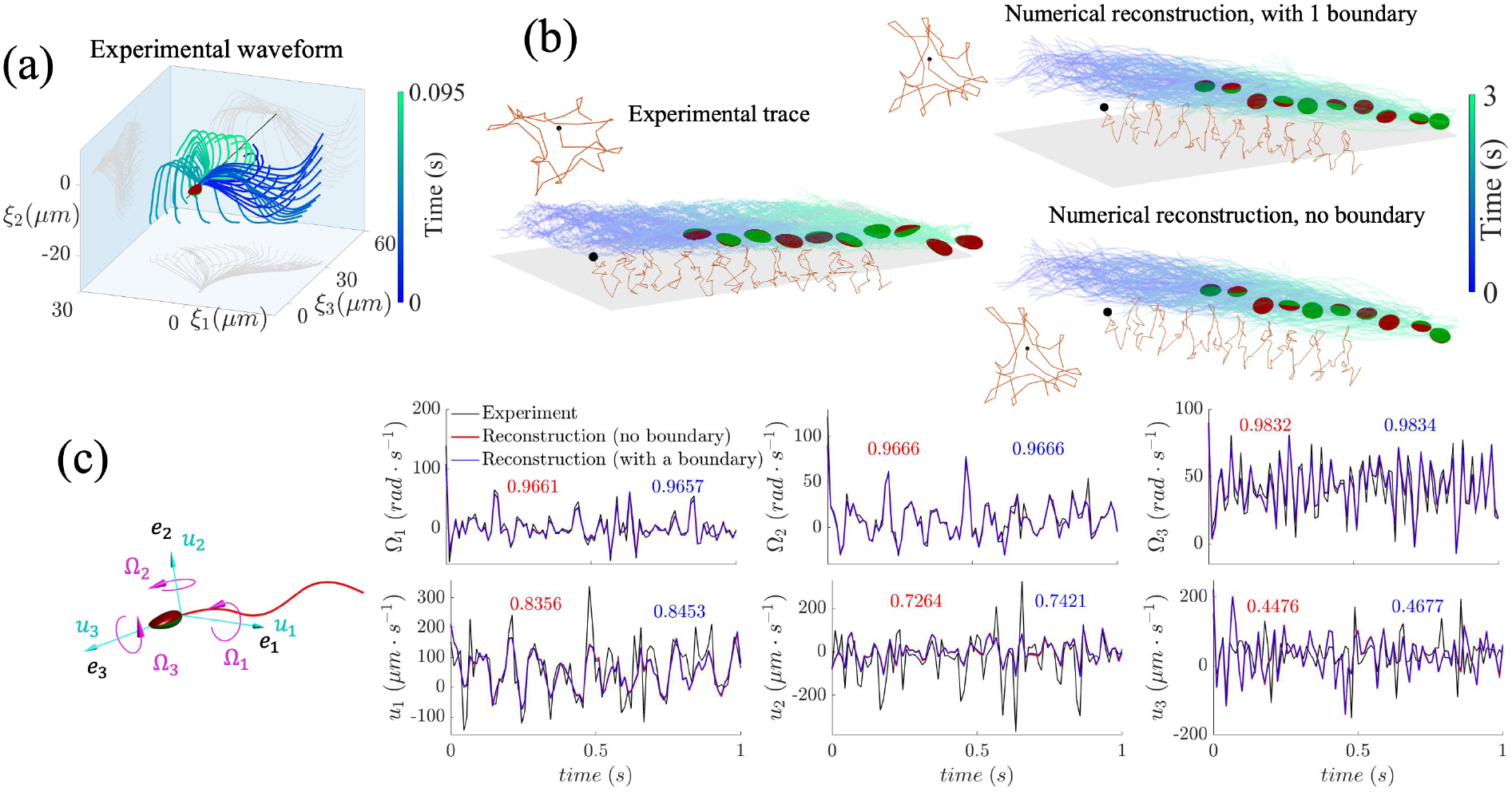
3D reconstruction for the observed human sperm. (a) Experimental waveform schematic in the body frame of reference. (b) Comparison of sperm traces between experimental (left) and reconstructed (right) results in the laboratory frame. The numerically reconstructed trace on the top takes boundary effects into consideration, with the initial distance from the sperm to the boundary equal to 0.2*L* (*L* for flagellum length), while the counterpart at the bottom presents sperm swimming in an unbounded fluid. Different colors are used to distinguish the upside and downside of the sperm head, red curves depict the trajectory shape of the mid-flagellar point (details shown in the insets, viewing from head to tail), and grey planes represent the boundary surfaces. (c) Comparison of sperm swimming speeds between experimental and reconstructed results. Sperm snapshot is superimposed to show the definition of the linear (*u*_1,2,3_) and angular (Ω_1,2,3_) speeds. The speeds for the reconstructions with and without a boundary are both provided, and the cross-correlation values for the 6 speeds between the experiment and the 2 reconstructions are given, in red and blue colors, respectively.

Numerical reconstructions on the 3D free sperm motility are performed using the experimental waveform, and the reconstructions are non-dimensionalized according to the measured flagellum length, *L* ≈ 59µ*m*, and waveform frequency, 10.3*Hz*. To model the sperm that swims immediately above the coverslip glass, we take boundary effects into consideration. Fig. 1(b), Fig. S1 and Supplemental Material, Videos 3 and 4 present the reconstructed sperm swimming in the lab frame, with different boundary conditions: no boundary and with boundary, whose distance from the sperm is 0.2*L*, 0.5*L* and 1*L*, respectively. Taking the experimentally observed sperm trajectory as the benchmark, the reconstruction results for the cases with and without a boundary all show excellent agreements.

To quantitatively validate our reconstructions, we measured the angular (linear) speeds, Ω_1,2,3_ (*u*_1,2,3_), around (along) the head basis vectors, ***e***_1,2,3_, from data in the lab frame, as defined in Fig. 1(c). The speeds for all simulations, shown in Fig. 1(c) and Fig. S2, are compatible with the experimental benchmark. The cross-correlation results between the experimental speeds and the counterparts of, for example, the no-boundary simulation are about 0.97, 0.97, 0.98, 0.84, 0.73, and 0.45, respectively, for Ω_1,2,3_ and *u*_1,2,3_. The consistency among all the reconstructions that incorporate or exclude a boundary implies that, for the experimentally reconstructed waveform, the influence of the hydrodynamics due to the presence of a boundary on sperm speeds and trajectory is marginal.

### 2.2 3D opposed vortices around human sperm

We investigate 3D hydrodynamics around the observed human sperm based on our verified reconstructions, and show the temporally averaged flow fields that comove with the head center in Fig. 2. The time-averaged fluid flows are obtained within the representative swimming cycles, the global CW revolutions shown in Fig. 1(b), and the dimensionless flow velocity is in units of flagellar length/ beat cycle. The flow structure in Fig. 2(a), where the boundary distance is 0.2*L*, shows disrupted circulations, while when the boundary distance increases to 0.5*L* and *L* (Fig. 2 (b) and (c)), the flow circulations become more organized. Finally, for the no-boundary case, Fig. 2(d) exhibits unperturbed 3D opposite vortices along the sperm body, due to the well-known sensitivity of the fluid flow due to a nearby wall.

**Figure 2.**
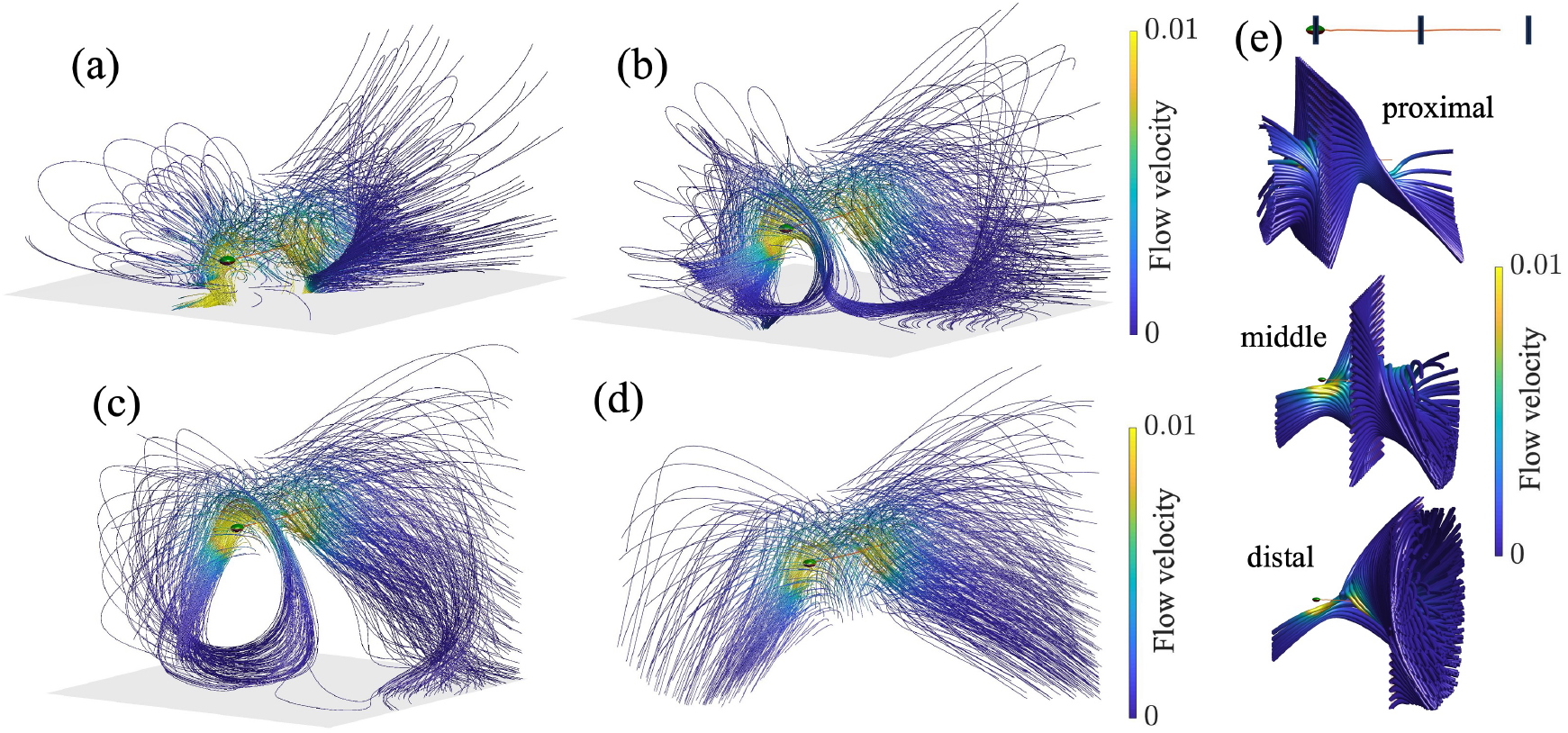
Time-averaged fluid flow around the reconstructed human sperm. (a) - (d) Average flows for the reconstructed sperm with different boundary conditions. From (a) to (c), perturbed opposite vortices, where grey planes represent the boundary, with boundary height 0.2*L*, 0.5*L* and *L*, respectively; (d) unperturbed opposite vortices, without a boundary. (e) 3D streamlines starting from different cross-sections of the sperm body, with section positions indicated by black vertical lines in the top inset. Flow velocity is visually represented through the colors of the streamlines.

Fig. 2(e) provides the details of the opposing vortices by illustrating the 3D streamlines starting from different flow regions. The fluid flow at the region proximal to the sperm head is dominated by CCW swirls when viewing from head to tail, while the flow near the distal flagellum tip is primarily featured by CW swirls. In the midflagellar region, CCW and CW vortices coexist. The streamlines in Fig. 2(e) first converge to the center of the corresponding cross sections, and then extend to one or both sides of the cross sections. Analogous near-field profiles have been described before for 2D flow predictions [11, 12, 19] or 3D flow imaging [8], showcasing the expulsion of fluid along the symmetry axis of the swimmer and the influx of fluid from the transverse direction. The flow convergence radii shown in Fig. 2(e) change along the sperm body, first decreasing and then increasing in the direction from head to tail, consistent with the predicted vortex patterns surrounding a swimming bacterium [20]. Similar swirls have been documented by [8], but are for tethered sperm, thus displaying unidirectional circulations. By comparison, the opposing vortices around a free sperm are a manifestation of the torque-free and force-free conditions on the free-swimming sperm dynamics [20–22]. Fig. 2 thus shows 3D near-field flow measurements around a freely swimming human sperm.

### 2.3 Virtual sperm models recapitulate the 3D experimental vortices

To explore the generality of the measured opposed vortices, 3D virtual sperm models are employed using the well-known elliptical helicoid waveform pattern [5, 23–26], as depicted in Fig. 3(a), and without boundary effects. Fig. S3 shows the lab frame trajectories of the virtual models, regulated by various waveform rotation amplitudes determined by *∝* (see Methods). Although the virtual waveform is different from the experimental flagellar beating pattern (Fig. 1(a)), their resultant lab frame trajectories demonstrate similar features, i.e. local CCW loops and global CW revolutions, as shown in Fig.1(b). When the waveform rotation amplitude (*∝*) increases, the color-encoded traces in Fig. S3 indicate a slower sperm swimming speed, as discussed in [26–28].

**Figure 3.**
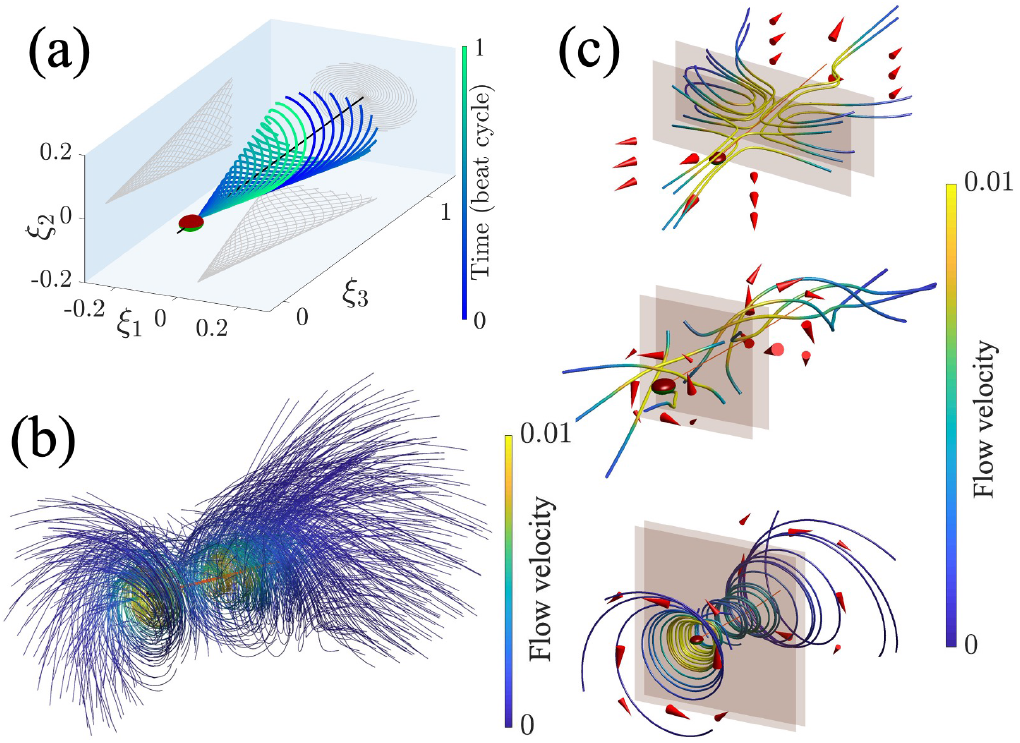
Time-averaged fluid flow around virtual sperm. (a) 3D virtual waveform in the body frame of reference. (b) Opposing vortices around the virtual sperm, with the waveform rotation amplitude *∝* = 0.6. (c) Average flows for the virtual sperm models, with *∝* = 0 (top), 0.4 (middle) and 1 (bottom). Brown planes indicate where the 3D streamlines start, red arrows denote the vortex direction, and flow velocity is visually represented through the colors of the streamlines.

The time-averaged fluid flow around the virtual sperm is shown in Fig. 3(b), characterized by CCW spirals in the front of the sperm and CW spirals in the rear. The close similarity of the near-field flow between the virtual model and reconstructed human sperm, despite the noticeable waveform disparity, suggests that the 3D opposite vortices represent a generic flow structure for a 3D free-swimming sperm in the bulk fluid. Besides, as shown in Fig. 3(c), such vortex pattern is intensified when the waveform rotation amplitude (*∝*) is large, accompanied with slower flow velocities. This may be responsible for the reduced velocity observed in the average flow generated by our 3D experimental waveform (Fig. 2) compared to that produced by the planar flagellar beating in [11]. In the far field, the average flow exhibits characteristics of a pusher type, as illustrated in Fig. S4, with flow velocity decaying as *∼ r*^−2^ [29–31].

### 2.4 Complexity of 3D instantaneous flow field

Fig. 4 (a) and (b) showcase the 3D instantaneous flow in the vicinity of our reconstructed human sperm, and the evolution of streamlines over time is shown in Supplemental Material, Video 5. The transient fluid flow changes markedly and rapidly, demonstrating strong dependence on time, compared with the slowly varying sperm movement, and the flow velocity is larger in magnitude than that of the time-averaged flow, agreeing with the predictions by [11, 14, 21]. In the far field, the instantaneous flow shown in Fig. S5 oscillates between pusher- and puller-type, in accordance with the conclusion drawn by [9, 11] for 2D waveforms.

**Figure 4.**
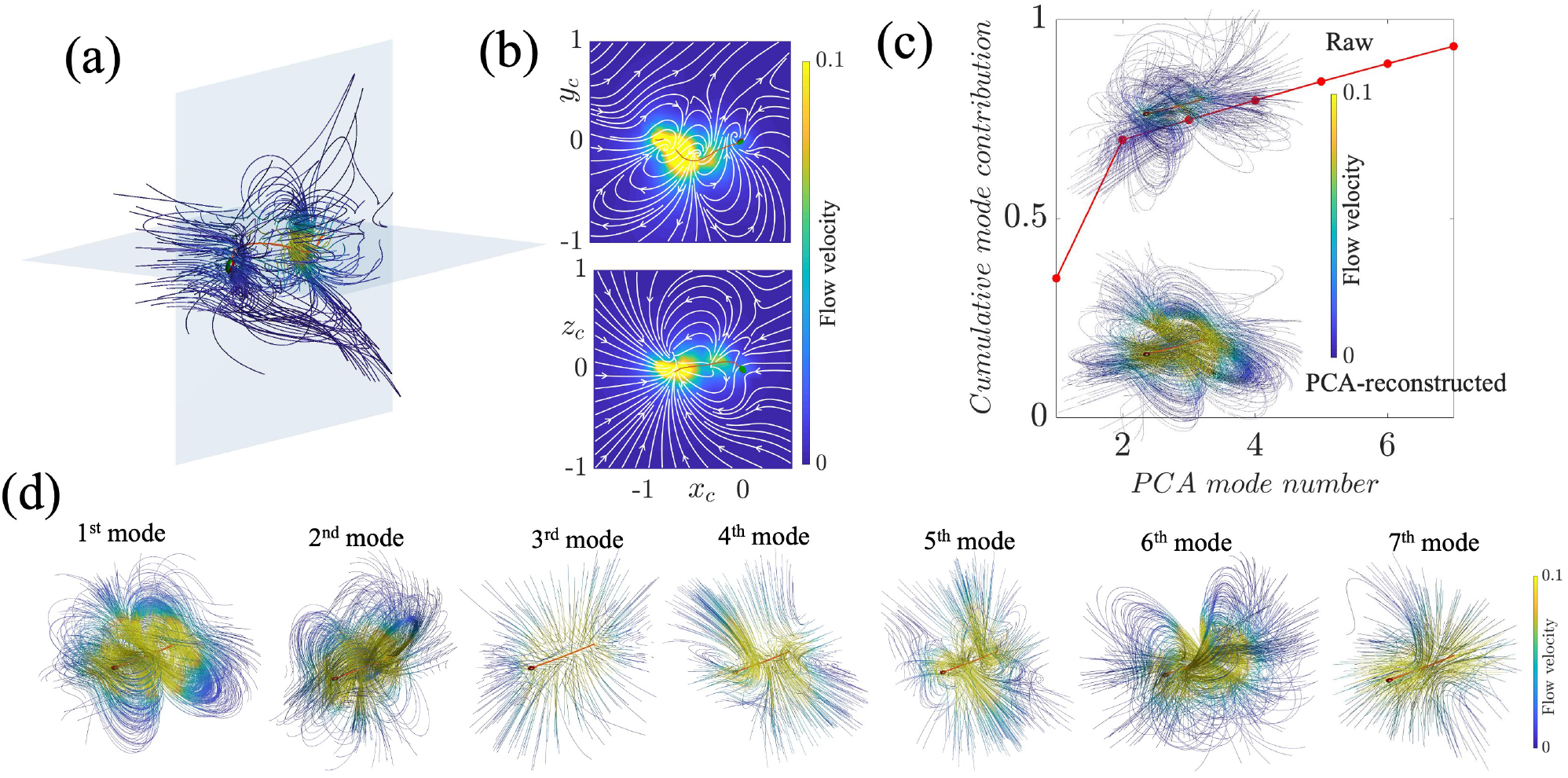
Instantaneous flow complexity. (a) 3D instantaneous flow around the reconstructed human sperm, where 2 blue planes indicate the XY and XZ planes for the 2D flow projections shown in (b). (c) PCA reconstruction of the instantaneous flow surrounding the virtual sperm (*∝* = 0.6): cumulative mode contribution for the first 7 PCA modes, with the raw instant flow and its PCA-reconstructed result displayed in the insets. (d) The first 7 modes employed in (c).

Principal component analysis (PCA) used to be applied to reduce the complexity of the time-varying flow induced by a planar flagellar beating [11], and here we employ this methodology to analyze the instant flow generated by 3D waveform. Given the remarkable resemblance in results between the human sperm and virtual model, we perform the PCA-decomposition based on the virtual model to circumvent experimental noise and investigate elementary mechanisms. Fig. 4(c) displays the raw transient flow and the corresponding PCA-reconstructed result, where the first 7 PCA modes are contained to capture 93.3% of the cumulative variance. The first and second PCA modes exhibit an approximate perpendicular relationship, as depicted in Fig. 4(d). This relationship is similarly found for the third (fifth) and fourth (sixth) modes, with the higher PCA modes possessing features of higher-order contributions.

## 3. Summary and conclusions

We report microscale fluid flow measurements of a free-swimming human sperm in low viscosity fluid powered by a 3D beating flagellum. For this, we experimentally measured the true beat of the sperm flagellum in 3D, as observed from the micron-sized sperm head that translates and rotates during cell swimming. This required high-resolution capture of both flagellum shape and head orientation in 3D. The experimental waveform was then used in conjunction with computational low Reynolds hydrodynamics to independently predict all sperm kinematics, and subsequently measure the fluid flow structures around a 3D beating human sperm.

Our hydrodynamic analysis of human sperm reveals key insights. We show that the flow field for a 3D free-swimming sperm in the bulk of a fluid features vortices with opposing circulation along the sperm cell. Vortices located at the front of the sperm swirl in a CCW direction, matching the head spinning direction, while those at the rear follow a CW direction. This average flow pattern is also recapitulated by idealised waveform, indicating the generality of this 3D flow structure. It is worth noting that our study focuses on sperm swimming in a low viscosity fluid [11, 32, 33], and future investigations will need to address the 3D hydrodynamics in high viscosity and physiological media [2].

We also examine the impact of a nearby boundary on sperm flagellum movements and resultant fluid flows. It is observed that, for the same experimental waveform imposed numerically, the presence of a boundary has negligible effects on sperm trajectories, as well as its linear and angular velocities for the duration of the experimental capture. However, the boundary significantly perturbs the 3D fluid flow structure when compared with the unbounded case. Near a wall, the average flow pattern shows perturbed vortical circulations. Moreover, our analysis of 3D instantaneous flow reveals its spatial and temporal complexity. The PCA decomposition for the 3D instantaneous flow suggests high-order contributions for multipolar expansion of singularities, as attempted for 2D beating patterns [11].

## 4 Methods

### 4.1 Sperm preparations

Sperm samples were obtained from healthy donors that went through a period of at least 48 hours of sexual abstinence. Donors were informed and asked to sign a written consent. The samples collected fulfilled the requirements established by the WHO. Readily after recollection, the samples went through a swim up separation for 1 hour with Ham’s F-10 medium at 37*°C* in a humidified atmosphere of 5% CO2 and 95% air. The physiological solution contained the following ingredients (expressed in mM): 94 NaCl, 4 KCl, 2 CaCl2, 1 MgCl2, 1 Na Piruvate, 25 NaHCO3, 5 Glucose, 30 HEPES, 10 Lactate at pH 7.4.

### 4.2 3D imaging microscopy

The experimental setup was resting on an optical table (TMC) to avoid external vibrations. A piezoelectric device (P-725 Physik Instruments, MA, USA) was mounted between an inverted microscope (Olympus IX71) and a 60X water objective lens (Olympus UIS-2 LUMPLFLN). A servo controller E-501 (Physik Instruments, MA, USA) coupled to a high current amplifier (Physik Instruments, MA, USA) was used to control the oscillations of the piezoelectric device. A symmetric saw signal was generated with a E-506 (Physik Instruments, MA, USA) with a frequency of 90*Hz* and amplitude of 20µ*m* to induce the oscillations in the piezoelectric device. A high speed camera NAC Q1v synchronized with the servo-controller recorded up to 3.5 seconds with a resolution of 640×480 and at 8000 fps. The cell samples were put in a chamber Chamlide (CM-B18-1) and the temperature was kept constant at 37*°C* with a thermal controller (Warner Instruments TCM/CL-100). A detailed description can be found in references [16, 34–36].

### 4.3 Experimental waveform acquisition

We record the 3D free swimming movement of human sperm in the laboratory frame of reference by capturing translational data of the flagellum motion, ***X***_*l*_ = (*x*_*l*_, *y*_*l*_, *z*_*l*_), and rotational information of head orientation, ***E*** = [***e***_1_, ***e***_2_, ***e***_3_]. As shown in Fig. 1(c), the head basis ***E*** is expanded by the unit vector ***e***_3_, which is along the head longitudinal axis and points forwards, ***e***_1_, normal to ***e***_3_ and lying in the flattening plane of the head, and ***e***_2_, which is orthogonal to the other two unit vectors by the right-hand rule. We first translate the experimental lab frame coordinates ***X***_*l*_ according to the position of the first flagellar point, and then rotate the flagellum shape through the orientation matrix ***E*** to obtain the flagellum beating relative to a fixed sperm head over time, namely waveform in the body frame of reference, referring to **ξ** = (ξ_1_, ξ_2_, ξ_3_) in Fig. 1(a).

The raw waveform is further reconstructed via PCA. The body frame flagellar data is mapped to a spatiotemporal matrix ***D***_*i*_*∝* = [ξ_1_(*t*_*i*_, *s*_*∝*_), ξ_2_(*t*_*i*_, *s*_*∝*_), ξ_3_(*t*_*i*_, *s*_*∝*_)], with time discretized into *n* values, *t*_1_, …, *t*_*n*_, and arc length discretized into *m* values, *s*_1_, …, *s*_*m*_. The temporally averaged result for each spatial point is denoted as 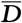 to form the covariance matrix 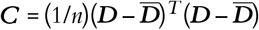, whose eigenvectors are known as PCA modes after being ordered by the size of the associated eigen-values. By projecting the raw flagellar data onto the span of the PCA modes, we can obtain the PCA-reconstructed waveform, 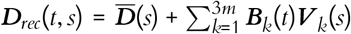, where ***B***_*k*_(*t*) is the temporal shape score for the corresponding mode ***V***_*k*_(*s*). We utilize the first 22 PCA modes to reconstruct the experimental waveform, capturing 99.91% of the cumulative variance. Furthermore, we employ the MATLAB functions “csaps” and “fnval” to smooth and interpolate the time-dependent coefficient ***B***_*k*_(*t*) for subsequent hydrodynamic calculations.

### 4.4 Hydrodynamic simulations for sperm swimming

We employ the framework of regularized Stokeslet method [37–39], using the nearest-neighbour discretization [18, 40, 41]developed by Smith’s group, to address the non-local hydrodynamics involved in sperm swimming. The flow velocity at a spatial point ***x*** in an unbounded fluid, driven by a regularized force of strength ***F*** at the location ***X***_*F*_, can be represented as 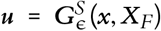 *·* ***F***, where 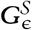 is the regularized Stokeslet. Sperm motility in a free space can be expressed using the boundary integral over the body surface ∂*D*, 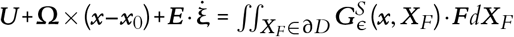, where ***U*** and **Ω** are the linear and angular velocities of the body frame relative to the lab frame, respectively, ***x***_0_ is the origin of the body frame, and the overdot of **ξ** denotes a time derivative of the body frame coordinate. The three unknown variables ***U*, Ω** and ***F*** can be calculated numerically by augmenting the mobility problem with force and torque balance equations, 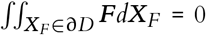 and 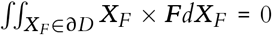, respectively. As such, lab frame kinematics of 3D free-swimming sperm can be predicted from a prescribed waveform in the body frame, as demonstrated in Fig. 1(b) and Fig. S3.

Apart from the sperm motility in an unbounded fluid, our models also consider boundary effects to mimic the cell that swims immediately above the cover glass. We utilize the regularized no-slip image system and replace the integral kernel with 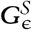, the regularized Blakelet that consists of a regularized Stokeslet and its mirror image, a Stokes-dipole and a potential dipole [42–45]. Once the sperm lab frame coordinates and Stokeslet force strength are determined, the surrounding flow velocity at ***x***_*f*_ can be calculated from the Stokeslets distributed over the sperm body, i.e. 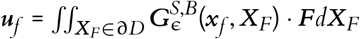. Fig. S6 presents the lab frame fluid flows for the sperm swimming near a wall, and compares the non-zero flow velocity in the sperm beating plane and the zero velocity on the boundary, thus validating our implementation of the no-slip boundary condition and the use of the image system. Both experimental and virtual waveforms are dimensionless, with time quantified in terms of beat cycles and the flagellum length normalized to 1, such that the dimensionless velocity is in units of flagellar length/ beat cycle. For the observed human sperm, the beat frequency is measured via fast Fourier transform.

### 4.5 Sperm linear and angular speed measurements

Sperm swimming speeds, **Ω** and ***u***, relative to the head basis vectors (***e***_1,2,3_) in the laboratory frame of reference are measured for quantitative comparisons between experimentally observed sperm motility and numerical reconstructions, referring to Fig. 1(c). The linear speed of the rigid sperm head, ***u*** = *u*_1_***e***_1_ + *u*_2_***e***_2_ + *u*_3_***e***_3_, is calculated by 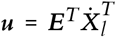, where head orientation ***E*** is already known and the time derivative, represented by the overdot, of head position ***X***_*l*_ can be readily obtained. The angular speed of sperm head and its orientation satisfy the relationship of *ė*_*i*_ = **Ω** *×* ***e***_*i*_, and thus we have **Ω** = (*ė*_2_ *·* ***e***_3_, *ė*_3_ *·* ***e***_1_, *ė*_1_ *·* ***e***_2_). Therefore, the angular speed **Ω** can be derived from the easily available head orientation and its time derivative.

### 4.6 Virtual sperm waveform

The beat envelope for our 3D virtual waveform **ξ**, as depicted in Fig. 3(a), has been widely described in the literature [5, 24–26], and can be prescribed as below:

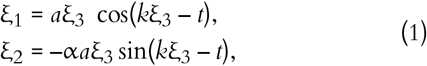

where *a* = 0.2 represents the modulating amplitude of the flagellar beating [26, 46], *k* is the wave number, taken as 2π according to the estimations from the observed waving patterns [8, 47, 48], and *∝* dictates the waveform rotation amplitude, ranging from 0 to 1 in this work, with the positive sign inducing a left-handed helicoid [5, 24, 49]. Here, we investigate the virtual sperm motility and its surrounding flow field by adjusting the 3D waveform component *∝*, see Fig. 3(c) and Fig. S3.

## 5. Data availability

The code, data and materials used in the analyses have been deposited in the public database https://github.com/polymaths-lab/3D-sperm-flow. Additional datasets related to this paper may be requested from the authors.

## 6 Acknowledgements

The authors acknowledge the computational facilities and team of the Advanced Computing Research Centre, University of Bristol: http://www.bristol.ac.uk/acrc/. Xiaomeng Ren acknowledges financial support of China Scholarship Council through Grant 202006830002. Dirección General de Asuntos del Personal Académico, Universidad Nacional Autónoma de México [DGAPA/UNAM; grants IN204922 (to A.D.) and IN105222 (to G.C.)]. P.H.-H. and A.D. acknowledge support from the Chan Zuckerberg Initiative (DAF grant 2020-225643 and 2023-329644) and NIH (RO1 HD038082-17A1), respectively.

## 7. Author information

### 7.1 Author contributions

X. R., H. B.-G., A. D., and G. C. designed the research. X. R. and H. B.-G. developed the main ideas, performed data analyses, and contributed to reviewing and editing the manuscript. X. R. performed mathematical data analysis and produced the initial manuscript. P. H.-H., and F. M. performed image processing. F. M., P. H.-H., and G. C. performed the experiment.

### 7.2 Corresponding author

Correspondence to Alberto Darszon, Gabriel Corkidi, and Hermes Bloomfield-Gadêlha.

